# Common ancestry of heterodimerizing TALE homeobox transcription factors across Metazoa and Archaeplastida

**DOI:** 10.1101/389700

**Authors:** Sunjoo Joo, Ming Hsiu Wang, Gary Lui, Jenny Lee, Andrew Barnas, Eunsoo Kim, Sebastian Sudek, Alexandra Z. Worden, Jae-Hyeok Lee

## Abstract

Homeobox transcription factors (TFs) in the TALE superclass are deeply embedded in the gene regulatory networks that orchestrate embryogenesis. Knotted-like homeobox (KNOX) TFs, homologous to animal MEIS, have been found to drive the haploid-to-diploid transition in both unicellular green algae and land plants via heterodimerization with other TALE superclass TFs, representing remarkable functional conservation of a developmental TF across lineages that diverged one billion years ago. To delineate the ancestry of TALE-TALE heterodimerization, we analyzed TALE endowment in the algal radiations of Archaeplastida, ancestral to land plants. Homeodomain phylogeny and bioinformatics analysis partitioned TALEs into two broad groups, KNOX and non-KNOX. Each group shares previously defined heterodimerization domains, plant KNOX-homology in the KNOX group and animal PBC-homology in the non-KNOX group, indicating their deep ancestry. Protein-protein interaction experiments showed that the TALEs in the two groups all participated in heterodimerization. These results indicate that the TF dyads consisting of KNOX/MEIS and PBC-containing TALEs must have evolved early in eukaryotic evolution, a likely function being to accurately execute the haploid-to-diploid transitions during sexual development.

**Author summary:** Complex multicellularity requires elaborate developmental mechanisms, often based on the versatility of heterodimeric transcription factor (TF) interactions. Highly conserved TALE-superclass homeobox TF networks in major eukaryotic lineages suggest deep ancestry of developmental mechanisms. Our results support the hypothesis that in early eukaryotes, the TALE heterodimeric configuration provided transcription-on switches via dimerization-dependent subcellular localization, ensuring execution of the haploid-to-diploid transition only when the gamete fusion is correctly executed between appropriate partner gametes, a system that then diversified in the several lineages that engage in complex multicellular organization.

## Introduction

The homeobox transcription factors (TFs) are ubiquitous in eukaryotes, carrying a DNA-binding homeodomain typically 60 amino acids, that folds into three α-helices [1]. The atypical or TALE (Three Amino acid Length Extension) superclass of homeobox TFs shares a three-amino-acid insertion between helix 1 and 2 and plays essential roles during embryonic development by participating in interactive TF networks. In animals, MEIS- and PBC-class TALE proteins, such as Meis/Hth and Pbx/Exd, form heterodimers that in turn form ternary complexes with HOX-class homeobox TFs, determining cellular fates along the anterior-posterior axis of the developing embryo [2,3]. In plants, the interacting KNOX- and BELL-class TFs in the TALE group play critical roles during organ formation and the vegetative-to-reproductive transition in the undifferentiated cell mass known as the shoot apical meristem [4,5].

The heterodimerization of TALE proteins serves as a trigger for precise execution of developmental programs. Prior to heterodimerization, animal PBX proteins are localized in the cytosol, and upon binding to MEIS, they translocate to the nucleus [6,7]. Similar heterodimerization-dependent translocation is also observed for KNOX-BELL pairs in the plant *Arabidopsis,* implying that this mechanism is a conserved regulatory feature of TALE proteins [8]. In addition, TALE proteins differ in their DNA-binding specificity [9,10], which is primarily determined by the homeodomain residues at positions 47, 50, and 54 [11], and heterodimerization increases target affinity by bringing two such DNA-binding domains together.

TALE-heterodimerization is mediated by class-specific homology domains located on the N-terminal side adjacent to the homeodomain [12,13]. Animal MEIS and plant KNOX class proteins share readily identifiable homology in their heterodimerization domain, leading to the proposal of an ancestral TALE class named MEINOX [12]. In contrast, their partner classes -- PBC and BELL -- exhibit no apparent homology in their heterodimerization domains. Short shared sequence motifs and common secondary structures have been found within the heterodimerization domains between MEINOX and PBC or BELL [14,15], but their extent of the homology requires adequate taxon sampling to recover ancestral relationships.

An ancestral functions of TALE-TALE heterodimerization was revealed in studies of the unicellular green alga *Chlamydomonas reinhardtii:* the KNOX ortholog GSM1 and a second TALE protein GSP1 form heterodimers immediately after the fusion of sexual gametes, and these drive the haploid-to-diploid transition by activating >200 diploid-specific genes and inactivating >100 haploid-specific genes [10,16,17]. In subsequent studies, plant-type TALE-TALE heterodimers between KNOX and BELL were shown to be required for the haploid-to-diploid transition of the moss *Physcomitrella patens* [18,19]. Given the conserved role of TALE heterodimerization as a developmental switch in the sexual life cycle of the plant lineage, understanding its origins and diversification promises to shed light on the evolution of developmental mechanisms during eukaryotic radiation and the emergence of land plants.

To delineate the ancestry of plant-type TALE heterodimerization, we performed a phylogenetic and bioinformatics analysis of TALE TFs in the three algal radiations of the Archaeplastida supergroup, the descendants of a single endosymbiosis event > one billion years ago [20,21]. Our analysis showed that the TALEs were already diversified into two groups at the origin of Archaeplastida, one sharing KNOX-homology and the other sharing PBC-homology. Together with our protein-protein interaction data, we propose that all TALE classes participate in heterodimerization networks via the KNOX- and PBC-homology domains between the two ancestral groups.

## Results

### TALEs in Archaeplastida are divided into two groups, KNOX and non-KNOX

The Archaeplastida consists of three monophyletic phyla [22,23] (Fig 1). 1) Viridiplantae include two divisions, Chlorophyta -- chlorophytes and prasinophytes (a paraphyletic group of seven lineages [24]) -- and Streptophyta -- charophyte algae and land plants [25]. 2) Rhodophyta (red algae) include diverse unicellular and multicellular organisms that diverge into four major lineages [26] (S1 Spreadsheet). 3) Glaucophyta members include only four cultured genera and possess plastids that carry ancestral features of the cyanobacterial symbiont that gave rise to photosynthetic organelles in eukaryotes [27].

**Fig 1.**
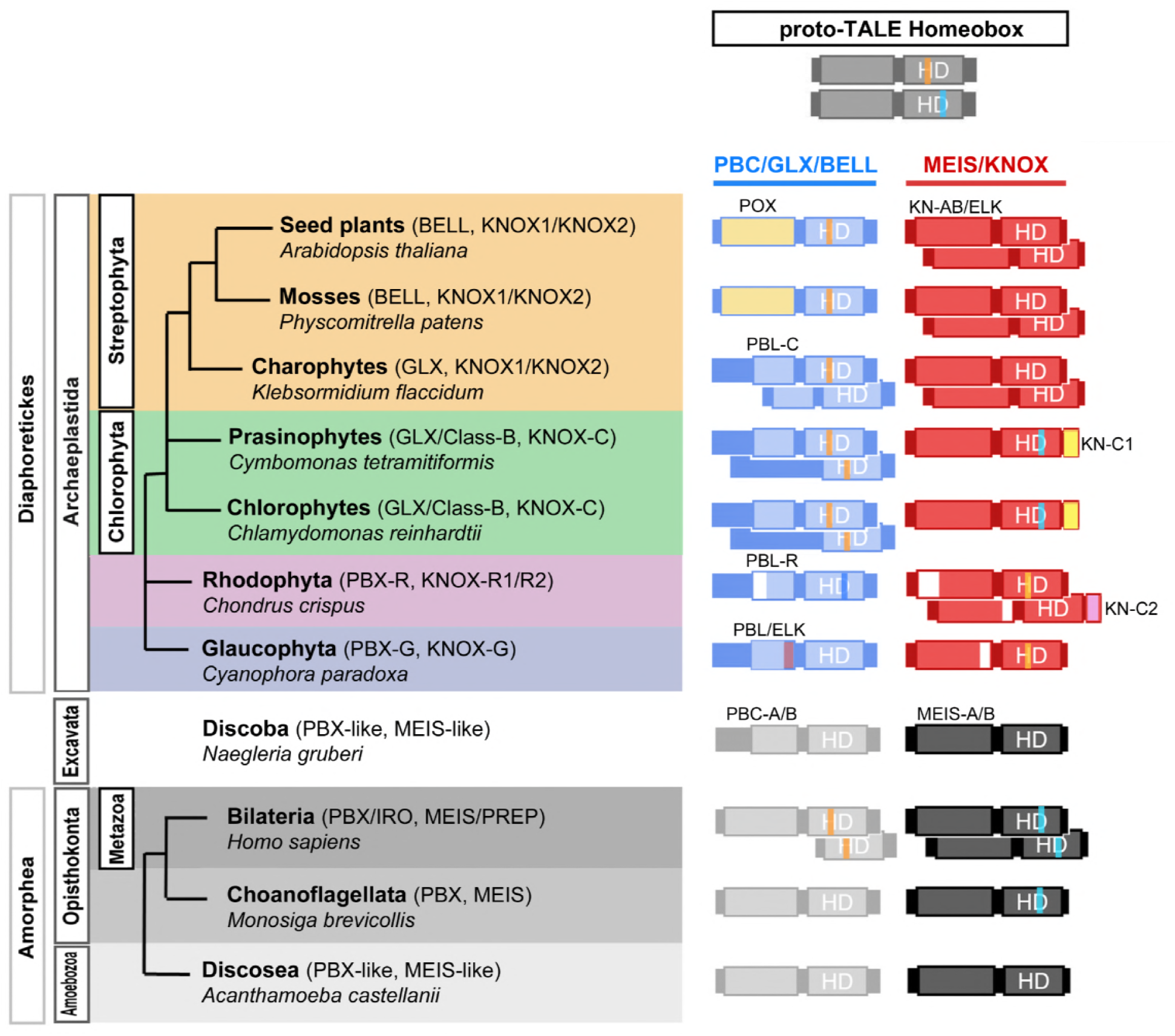
Common origin of heterodimerizing TALE homeobox TFs. We propose that a homodimerizing proto-TALE protein duplicated (top) prior to the major bifurcation resulting in animals/fungi/amoebae vs. algae/plants. Lineage-specific diversification soon followed, generating heterodimeric configurations distinct at the phylum-level. These configurations usually couple members of the PBC/PBX/GLX group that shares PBC-homology domains and MEIS/KNOX group that shows homology in the KN-A/B domains N-terminal to the homeodomain. Each lineage possesses one or two classes of potential heterodimeric partners. Major TALE classes are mapped onto the eukaryotic phylogeny. A representative species name is given for each analyzed lineage. Open boxes in the domain diagrams indicate the absence of MEINOX for PBX-Red, KN-A for KNOX-Red1 and ELK for KNOX-Red2. Colored vertical lines in the HD indicate two proposed ancestral introns at 44/45 (orange over ‘H’ In HD) and 48[2/3] (blue over ‘D’ in HD), whose alternating existence between the two groups suggests independent diversification of TALE heterodimerization.

To collect all the available homeobox protein sequences, we performed BLAST and Pfam-motif searches against non-plant genomes and transcriptome assemblies throughout the Archaeplastida (S1 Spreadsheet), identifying 327 proteins from 55 species as the Archaeplastida homeobox collection (29 genomes and 18 transcriptomes; S2 Spreadsheet). Of these, 102 possessed the defining feature of TALE proteins, a three-amino-acid insertion between aa positions 23-24 in the homeodomain [28]. At least two TALE genes were detected in most genomes except five genomes in the Trebouxiophyceae class of the Chlorophyta (S1 Spreadsheet; see S1A Notes for further discussion of the absence of TALEs in Trebouxiophyceae).

The collected TALE sequences were then classified by their homeodomain features using a phylogenetic approach, with TALEs from animals, plants, and early-diverging eukaryotes (Amoebozoa and Excavata) as outgroups (S1 Fig). The resultant TALE homeodomain phylogeny distinguished two groups in all three phyla of Archaeplastida (Fig 2). 1) The KNOX-group as a well-supported clade displayed a phylum-specific cladogram: two Glaucophyta sequences at the base (as KNOX-Glauco) were separate from the next clade, which combines Rhodophyta sequences (as KNOX-Red1) and a Viridiplantae-specific clade with strong support (92/90/1.00). 2) The non-KNOX group, including the BELL and GSP1 homologs, contained clades of mixed taxonomic affiliations. These analyses showed that the TALE proteins had already diverged into two groups before the evolution of the Archaeplastida and that the KNOX-group is highly conserved throughout Archaeplastida.

**Fig 2.**
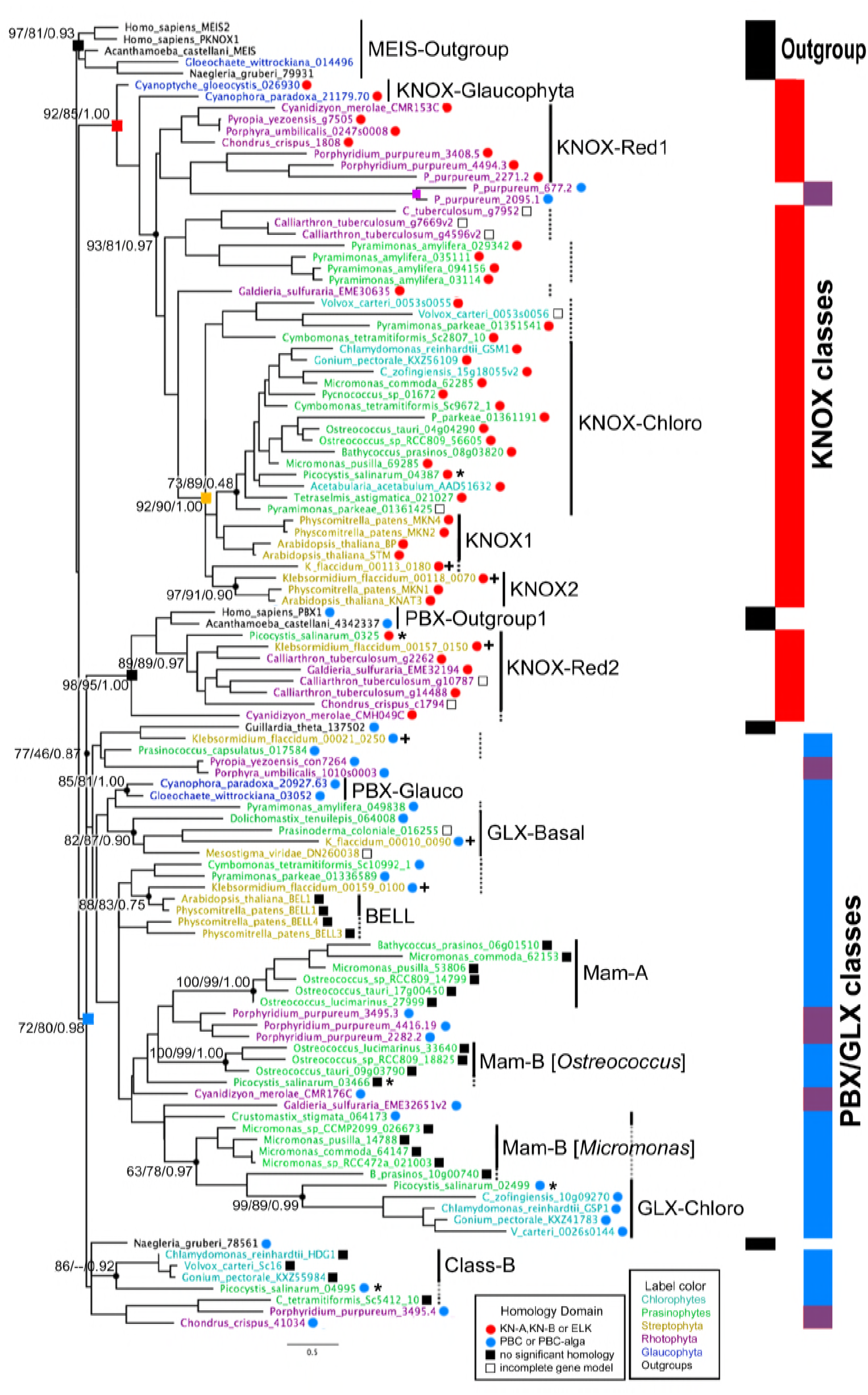
Maximum likelihood (ML) phylogeny of the TALE superclass homeodomain in Archaeplastida supports ancient division between KNOX- and non-KNOX TALE groups. The consensus tree out of 1000 bootstrap trees is shown. The three numbers shown at nodes are %bootstrap, %SH, and Bayesian posterior probability in support of clades. The tree contains two outgroup clades marked by black squares, and two Archaeplastida clades, one combining most KNOX sequences marked by the red square and the other combining all non-KNOX sequences marked by the blue square. Vertical bars on the right depict the distribution of outgroup, KNOX, and non-KNOX sequences. KNOX sequences are marked with red dots indicating the presence of KN-A or KN-B domains. GLX/PBX sequences are marked with blue dots indicating the presence of a PBC-homology domain. Truncated sequences not available for homology domain analysis are marked with open black boxes. Filled box indicates the absence of a KN-A/B or PBC-homology domain. Proposed classification is shown by black vertical lines. Dotted lines indicate sequences related to a class but placed outside the main clade for the class. PBX-Red sequences are found in four paraphyletic clades, marked by purple shades on the blue vertical bar. Sequence IDs containing the species name are colored by their phylogeny: Blue for Glaucophyta, purple for Rhodophyta, green for prasinophytes, light blue for the chlorophytes, orange for Streptophyta, and black for outgroups. The ruler shows genetic distance. All the sequences and their phylogenetic information are found in S2 Spreadsheet. *Gloeochaete_wittrockiana_014496 is considered as a sequence from a bannelid-type amoeba that contaminated the original culture (SAG46.84) for the MMETSP1089 transcriptome. **Association of KNOX-Red2 class sequences to Amorphea PBC sequences is attributed to a shared WFGN motif determining DNA-binding specificity of the homeodomain via convergent evolution.

### KNOX group sequences share the same heterodimerization domains throughout Archaeplastida

The next question was whether the plant KNOX class originated prior to the Viridiplantae phylum. The plant KNOX proteins and the Chlorophyta GSM1 possess KNOX-homology sequences, consisting of KN-A, KN-B and ELK domains, required for their heterodimerization with other TALE proteins [10]; therefore, the presence of the KNOX homology sequences suggest functional homology to the plant KNOX class. To collect homology domains without prior information, we performed ad-hoc homology domain searches among the KNOX group sequences. Using the identified homology domains as anchors, we carefully curated an alignment of the KNOX-group sequences combined with any other TALE sequences with a KNOX-homology, (S2 Fig). From this KNOX alignment, we defined KNOX-homologs as having amino acid similarity scores >50% for at least two of the three domains comprising the KNOX-homology region (S3 Spreadsheet for calculated domain homology). Using this criterion, all KNOX group sequences (excluding partial sequences) possessed the KNOX homology (Fig 2, marked by red dots following their IDs), indicating that the KNOX-homolog already existed before the evolution of eukaryotic photosynthesis as represented by the Archaeplastida.

In addition to the KNOX-homology, the same search also revealed two novel domains at the C-terminus of the homeodomain (S2 Fig): the first (KN-C1) was shared among the Chlorophyta sequences, and the second (KN-C2) was shared among a group of KNOX homologs in a clade outside the KNOX-group (KNOX-Red2).

### KNOX classes diverged independently among the algal phyla

In Viridiplantae, we found a single KNOX homolog in most Chlorophyta species, whereas KNOX1 and KNOX2 divergence was evident in the Streptophyta division, including the charophyte *Klebsormidium flaccidum* and land plants (Fig 2). The newly discovered KN-C1 domain was specific to the Chlorophyta KNOX sequences and found in all but one species *(Pyramimonas amylifera).* The absence of similarity between KN-C1 and the C-terminal extensions of KNOX1/KNOX2 sequences suggests independent, lineage-specific KNOX evolution in the Chlorophyta and Streptophyta (S2 Fig). We, therefore, refer to the Chlorophyta KNOX classes as KNOX-Chloro in contrast to the KNOX1 and KNOX2 classes in the Streptophyta.

The KNOX homologs in the Rhodophyta were divided into two classes: a paraphyletic group close to the KNOX-Chloro clade, named KNOX-Red1, and a second group near the PBX-Outgroup, named KNOX-Red2. KNOX-Red1 lacked a KN-A, whereas KNOX-Red2 lacked an ELK and shared a KN-C2 domain (S2 Fig). We consider KNOX-Red1 as the ancestral type, since the KNOX-Red1 sequences were found in all examined Rhodophyta taxa, whereas the KNOX-Red2 sequences were restricted to two taxonomic classes (Cyanidiophyceae and Florideophyceae). Interestingly, the KNOX-Red2 clade included two green algal sequences, with strong statistical support (89/89/0.97; Fig 2); these possessed a KN-C2 domain, suggesting their ancestry within the KNOX-Red2 class (S2 Fig; see S1B Notes for further discussion about their possible origin via horizontal gene transfer).

Available TALE sequences were limited for the Glaucophyta. We found a single KNOX homolog in two species, which possessed KN-A and KN-B domains but lacked an ELK domain. We termed these KNOX-Glauco.

### Non-KNOX group TALEs possess animal type PBC-homology domain, suggesting a shared ancestry between Archaeplastida and Metazoa

Following the identification of KNOX homologs, the non-KNOX group in the Archaeplastida was redefined as lacking KN-A and KN-B domains. Further classification of the non-KNOX group was challenging due to its highly divergent homeodomain sequences. However, we noticed that the number of non-KNOX genes per species was largely invariable: one in most Rhodophyta and Glaucophyta genomes and two in the majority of Chlorophyta genomes, suggesting their conservation within each radiation.

Our ad-hoc homology search provided critical information for non-KNOX classification, identifying a homology domain shared among all Glaucophyta and Rhodophyta non-KNOX sequences (Fig 3A and 3B). Since this domain showed a similarity to the second half of the animal PBC-B domain (Pfam ID: PF03792) known as heterodimerization domain [12], we named this domain PBL (PBC-B Like). Accordingly, we classified all the non-KNOX TALEs in Glaucophyta and Rhodophyta as a single PBC-related homeobox class, PBX-Glauco or PBX-Red. PBX-Glauco sequences also possessed the MEIONOX motif, conserved in the animal PBC-B domain, indicating common ancestry of PBC-B and PBL domains (Fig 3A).

**Fig 3.**
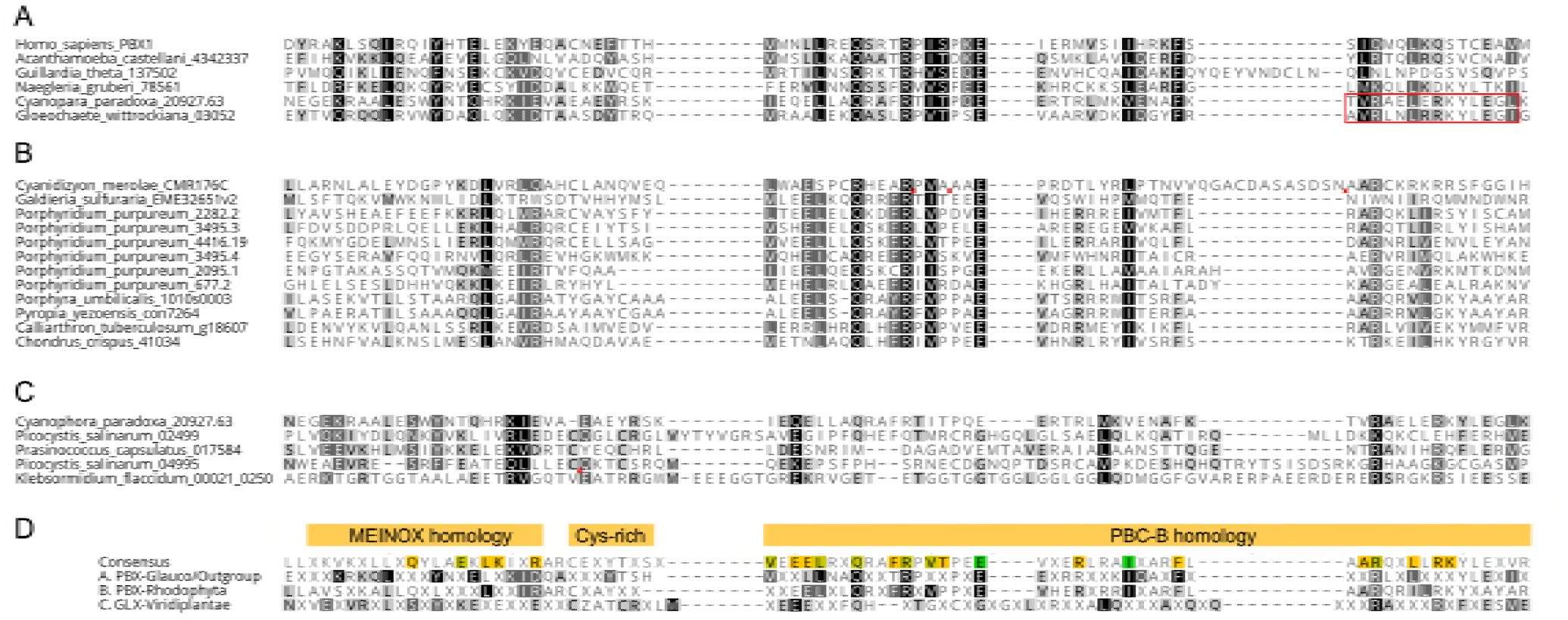
Archaeplastida Non-KNOX group TALEs possess a PBL domain sharing homology with metazoan PBC class TALEs. (A) PBL-Glauco domain alignment. Two Glaucophyta non-KNOX sequences possess a PBC-homology domain spanning MEINOX and C-terminal half of the PBC-B domains which is shared among the three outgroup TALE sequences analyzed in this study. (B) PBL-Red domain alignment. All Rhodophyta non-KNOX sequences possess a PBC-homology domain that can be aligned to the MEINOX/PBC-C domains. (C) PBL-Chloro domain. Four non-KNOX sequences show >10% amino acid identity to one of the other PBC-homology blocks presented in (A) and (B). Picocystis_salinarum_ü2499 is a founding member of GLX class with a PBL-Chloro domain. D. Comparison among PBC-homology domains. The top row shows the consensus made from the alignment of (A), (B), and (C) combined and the lower consensus sequences are collected from the individual alignments presented in (A), (B), and GLX alignment (S3 Fig). Amino acid letters in black with Gray shades, in white with light shades, and in white with black shades show more than 60%, 80%, or 100% similarity in each column.

### GSP1 shares distant PBC-homology together with other non-KNOX group sequences in Viridiplantae

A remaining question was the evolution of the Chlorophyta non-KNOX sequences that apparently lacked a PBC-homology. To uncover even a distant homology, we compared the newly defined PBL domains with the Chlorophyta sequences by BLAST (cut-off E-value of 1E-1) and multiple sequence alignments. This query collected three prasinophyte and one charophyte TALE sequences that possessed a MEINOX motif and a putative PBL-domain; however, they showed very low sequence identity among themselves (Fig 3C). Further query utilizing these four sequences identified 11 additional non-KNOX sequences. Nine of these were made into two alignments, one including GSP1 homologs and the other combining most prasinophyte sequences (S3A and S3B Fig). The two remaining sequences (Picocystis_salinarum_04995 and Klebsormidium_flaccidum_00021_0250) showed a homology to a PBX-Red sequence of *Chondrus cruentum* (ID:41034) in a ~ 200 aa-long extension beyond the PBL domain, suggesting their PBX-Red ancestry (another potential case of horizontal transfers; S4 Fig). All the Chlorophyta non-KNOX sequences that carry the PBL-homology domains were classified as GLX (GSP1-like homeobox) in recognition of the GSP1 protein of *Chlamydomonas* as the first characterized member of this class [29].

### Two non-KNOX paralogs of Chlorophyta heterodimerize with the KNOX homologs

Even with our sensitive iterative homology search, we could not identify a PBC/PBL-homology in about half of the Chlorophyta non-KNOX sequences. Since most Chlorophyta genomes possess one GLX homolog and one non-KNOX sequence without the PBL-homology domain, we refer the latter collectively to Class-B (S5 Fig). Exceptions were found in one prasinophyte clade (class Mamiellophyceae), whose six high-quality genomes all contain two non-KNOX sequences lacking the PBL-homology. Nonetheless, these non-KNOX sequences formed two groups, one more conserved and the other less conserved, referred to the Mam-A and Mam-B classes, respectively (S6 and S7 Fig). Considering the reductive genome evolution of the Mamiellophyceae [30], the conserved Mam-A class may be derived from an ancestral GLX class.

Two divergent non-KNOX classes in Chlorophyta led to a critical question about their dyadic networks. Previously studies had shown that TALE heterodimers required interaction between MEIS and PBC domains in animals and between KNOX and PBL domains in *Chlamydomonas* [6,10]. It was, therefore, predicted that all Glaucophyta and Rhodophyta TALEs form heterodimers via their KNOX- and PBL-homology domains. On the other hand, it remained to be tested whether the Chlorophyta TALEs lacking a PBL-domain can form heterodimers with other TALEs.

To characterize interaction network of TALE class proteins in Chlorophyta, we selected three prasinophyte species for protein-protein interaction assays: two species containing Mam-A and Mam-B genes *(Micromonas commoda* and *Ostreococcus tauri),* and another species *(Picocystis salinarum),* whose transcriptome contained one GLX and one Class-B sequence. In all three species, we found that KNOX homologs interacted with all examined non-KNOX proteins in Mam-A, Mam-B, Class-B, and GLX class (Fig 4A-4C). No interaction was observed between the two non-KNOX proteins in any of the three species (Fig 4A-4C). Similar to the GLX-KNOX heterodimerization, Mam-A and Mam-B also required additional domains outside the homeodomain for their heterodimerization with the KNOX homologs (S8 Fig). These results showed that the all divergent non-KNOX TALEs maintained their original activity to form heterodimers with the KNOX homologs. Observed interacting network among the TALE sequences is summarized in S9A Fig.

**Fig 4.**
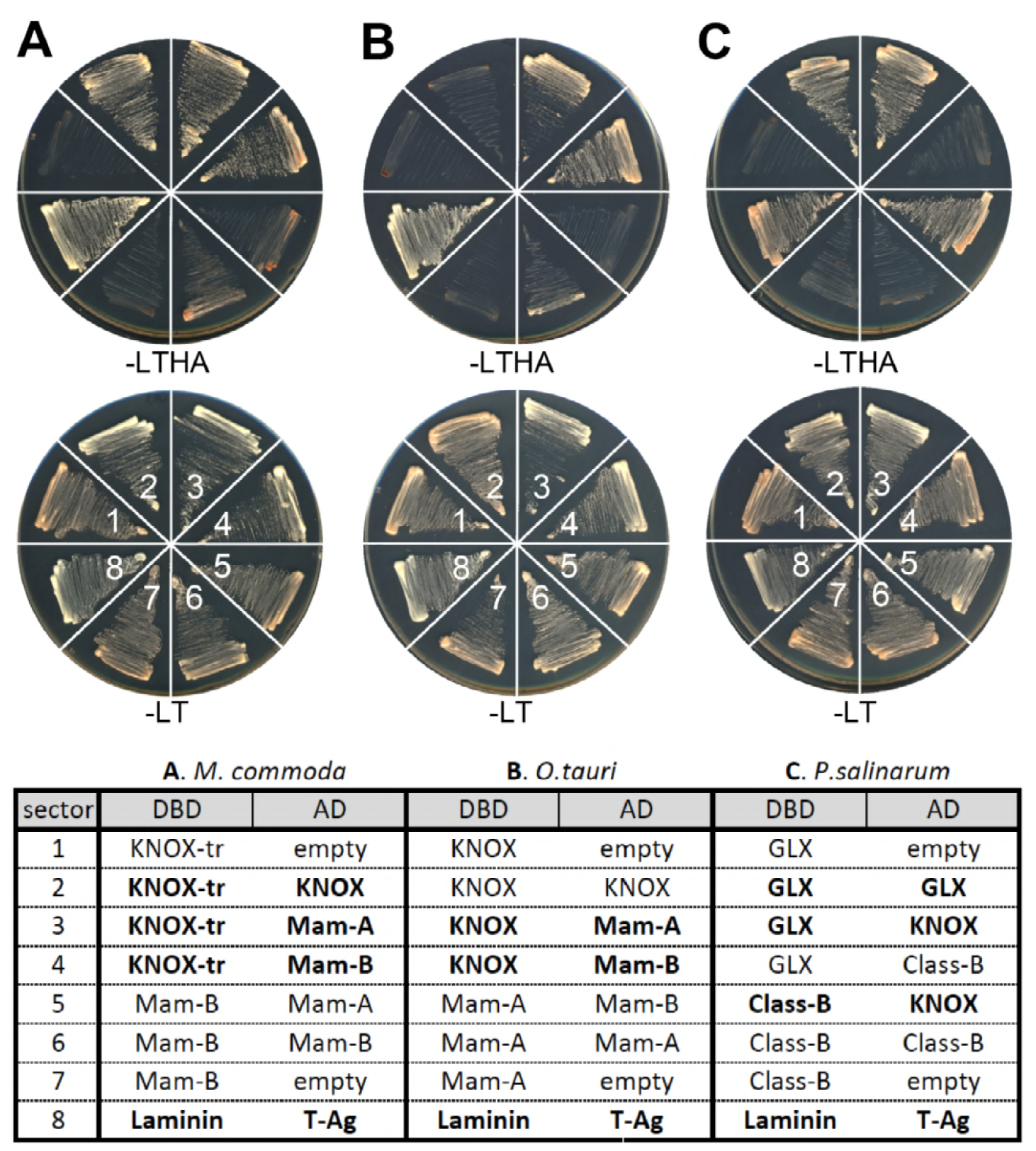
All Chlorophyta TALE TFs engage in heterodimerization networks. The bait constructs conjugated to the GAL4 DNA-binding domain and the prey constructs conjugated to the GAL4 transcriptional activation domain are listed in the table. Construct combinations, numbered 1-8, are arranged in wedges clock-wise, starting at 9 o’clock as labeled in (A). Interacting pairs confer yeast growth in Leu-/Trp-/His-/Ade- (-LTHA) medium. Confirmed interacting pairs are shown in bold faces in the table. The laminin and T-Antigen (T-Ag) pair, known to be interacting partners, was plated in the 8th sector as a positive control. (A) Assays using *M. commoda* TALEs. (B) Assays using *O. tauri* TALEs. C. Assays using *P. salinarum TALEs.* Class-A refers to the GLX-Chloro homolog. Details of the construct information are found in S5 Spreadsheet.

### TALE heterodimerization evolved early in eukaryotic history

Our discovery of the PBC-homology in Archaeplastida suggests common ancestry of the heterodimerizing TALES between Metazoa and Archaeplastida. It also predicted that other eukaryotic lineages might possess TALEs with the PBC-homology. Outside animals, the Pfam database contains only two PBC-B domain-harboring sequences, one from a Cryptophyta species *(Guillardia theta,* ID_137502) and the other from an Amoebozoa species *(Acanthamoeba castillian,* ID:XP_004342337)[31]. We further examined the Excavata group, near to the posited root of eukaryotic phylogeny [22]. A search of two genomes *(Naegleria gruberi* and *Bodo saltans)* collected 12 TALE homeobox sequences in *N.gruberi*, and none in *B.saltans*, of which we found one with a PBC-homology domain (ID:78561, Fig 3A) and one with a MEIS/KNOX-homology (ID:79931, S2 Fig). Our data suggest that the heterodimerization domains -- the PBC-homology and MEIS/KNOX-homology -- originated early in eukaryotic evolution and persisted throughout the major eukaryotic radiations.

### Intron-retention supports the parallel evolution of the heterodimeric TALE classes during eukaryotic radiations

The ubiquitous presence of dyadic TALEs raised next question: Are all the dyadic TALEs reported in this study the descendants of a single ancestral dyad, or do they result from lineage-specific evolution from a single prototypical TALE (proto-TALE) that does not engage in heterodimerization. To probe deep ancestry, we examined intron-retention, this being regarded as a long-preserved character and less prone to occur by homoplasy (a character displayed by a set of species but not present in their common ancestor) [32]. Five intron positions were shared by at least two TALE classes, of which the 44/45 and 48[2/3] introns qualified as the most ancestral since they were found throughout the Archaeplastida and Metazoa (S10 Fig).

The 44/45 and 48[2/3] introns showed an intriguing exclusive distribution between the two dyadic partners of each phylum: one possesses the 44/45 and the other the 48[2/3] intron (S10 Fig). This mutually exclusive pattern suggested that two TALE genes with distinct intron positions existed at the onset of the eukaryotic radiation. We consider the 44/45 intron position as the most ancestral, given that it was conserved in most non-TALE homeobox genes [12]. In this regard, we speculate that acquisition of the 48[2/3], and loss of the 44/45 intron, accompanied an early event wherein the proto-TALE with the 44/45 intron was duplicated to generate a second TALE with the 48[2/3] intron. Since the two intron positions were found within both the MEIS/KNOX and PBC/PBX/GLX groups, we propose that the duplicated TALEs arose early and diversified to establish lineage-specific heterodimeric configurations during eukaryotic radiations.

Given that the heterodimeric TALEs evolved in a lineage-specific manner, we asked what the proto-TALE looked like at the time it underwent duplication. The following observations suggest that the proto-TALE was a homodimerizing protein. First, the PBC-homology domains of PBX/GLX class proteins identified in the Archaeplastida includes the MEINOX-motif that was originally defined for its similarity to the MEIS/KNOX-homology domains (Fig 3) [14]. Second, PBX-Glauco sequences possess the ELK-homology within their PBL domain (Fig 3), which align well to the ELK domains of KNOX class sequences in Viridiplantae (S11 Fig). Therefore, the MEINOX-motif and ELK-homology across the heterodimerizing KNOX and PBX groups supported the common origin of heterodimerizing TALE groups from a single TALE by duplication followed by subfunctionalization.

## Discussion

### TALE endowment in Archaeplastida

Our study shows that all three Archaeplastida phyla possess TALEs, diverged into two groups with distinct heterodimerization domains, the KNOX group with KN-A/KN-B domains and the PBX (or GLX) group with PBL domains. The similarity between the KNOX/PBX and the animal MEIS/PBC dyads led us to identify homologous heterodimerization domains in the TALEs of other eukaryotic lineages including Excavata. Based on our findings, we hypothesize that the TALE heterodimerization arose very early in eukaryotic evolution.

During > 1 BY of Archaeplastida history, TALE TF networks have undergone three duplication events compared to the simple dyadic TALEs in Glaucophyta. In Viridiplantae, the KNOX class persists as a single member throughout the mostly unicellular Chlorophyta, whereas it duplicated into KNOX1 and KNOX2 in the multicellular Streptophyta [33]. In Rhodophyta, two KNOX classes, KNOX-Red1 and KNOX-Red2 differ in KN-A and KN-B domains, suggesting sub-functionalization. The third duplication event occurred in the non-KNOX group of the Chlorophyta, whose sequences then underwent rapid divergence in their homeodomain and heterodimerization domains, rendering their classification trickier than other classes. Despite this divergence, proteins in one of the two radiations (Class-B and Mam-B) were found to heterodimerize with KNOX homologs, suggesting that these non-KNOX members serve as regulators of KNOX/GLX heterodimers. We summarize our finding in Fig 1, S10B Fig.

### Is the plant BELL class homologous to the Chlorophyta GLX class?

The BELL class is the only non-KNOX class in land plants, sharing a POX (Pre-homeobox) domain (PF07526) [13] and lacking an identifiable PBL domain. The *K. flaccidum* genome, the only genome available in the charophyte lineage from which land plant emerged, contained three non-KNOX sequences, all possessing a PBL domain (Fig 3, S3,S4 Fig). Therefore, the lack of PBL-homology in the plant BELL class appears to be due to divergence or domain loss from an old charophyte class that had PBL-homology. We found an intron at the 24[2/3] homeodomain position of a *K. flaccidum* GLX homolog, which was previously identified as being specific to the plant BELL class (S8A Fig) [12], suggesting that the plant BELL class evolved from an ancestral GLX gene. More taxon sampling in charophytes is needed to confirm this inference.

### What would have been the critical drivers of TALE heterodimerization networks emerging from ancestral homodimers?

We found two conserved intron positions and shared sequence motifs between the KNOX- and PBX-groups, generating our hypothesis that a proto-TALE protein initially engaged in homodimerization and then duplicated and diversified into two heterodimerizing classes (Fig 1, S9A Fig). Heterodimerization-dependent subcellular localization [10,34], coupled with numerous combinations of distinct DNA-binding modules that fine-tune target specificity, then generated customized transcription-on switches.

During sexual development, it is critical to accurately detect the fusion of two cells before initiating diploid development and to make sure that the mating combines correct partner gametes. TF heterodimerization can implement both steps if one TF partner is contributed by each gamete. In fact, TALE heterodimerization plays a central role as a developmental switch for the haploid-to-diploid transition in green algae and land plants [10,19]. A similar haploid-to-diploid transition triggered by TF heterodimerization has recently been documented in *Dictyostelium* [35] and is well described in Basidiomycete fungi that utilize non-TALE homeobox proteins such as bW and bE [36,37].

Discovery of new prokaryotic life forms, especially in the Archaea domain, suggests that multiple symbiotic mergers of different life forms evolved into the proto-eukaryotes, possibly first as a symbiotic community, which then evolved into the last eukaryotic common ancestors (LECA) that rapidly diverged into the eukaryotic supergroups [38–40]. This eukaryogenesis model predicts that the proto-eukaryotes → LECA transition required the faithful transmission of traits between progenitor cells and their progeny to evolve as individual lineages by Darwinian selection. Under this hypothesis, we anticipate that the generation of the LECA may have been driven by the sexual mechanisms that distinguish a cellular merger between the common descendants from a merger between unrelated community members. Our proposal for the evolution of heterodimeric TALEs from the homodimeric proto-TALE may provide one of the necessary mechanisms for the first sexual mode of reproduction that might have driven the generation of the LECA from its proto-eukaryotic ancestors.

### Does expansion of heterodimerizing TALE TFs relate to the emergence of multicellular complexity?

Plant studies have shown that the duplicated KNOX classes serve distinct functions: the plant KNOX1 class regulates the differentiation of an undifferentiated cell mass into spores in mosses or leafy organs in vascular plants, and the plant KNOX2 class regulates the transition from haploid gametophytes to diploid sporophytes in mosses and controls secondary cell wall development in vascular plants [18,41–43]. We propose that the duplicated TALE heterodimers in the Streptophyta allowed independent regulation of cellular differentiation and life cycle transitions, priming the emergence of land plants by expanding the diploid phase of their life cycle from a dormant zygospore to a multicellular individual bearing many meiotic spores. The repertoire of TALE heterodimers continued to expand during land plant evolution, serving all the major organ differentiation programs in the diploid phase of their life cycle.

Can a similar expansion of TALE heterodimers be found during Metazoan evolution? Our search for TALE TFs in unicellular relatives of the Metazoa -- Spingoeca and Monosiga – revealed a simple configuration with one MEIS- and one PBC-like TALE (S12, S13 Fig), whereas at the Metazoan base one finds at least three MEIS-related classes and two PBC-related classes [44]. These findings suggest the occurrence of a similar expansion of a founding dyad during Metazoan evolution. Therefore, in both plants and animals, the TALE TF network seems to be redeployed for complex multicellularity, departing from its posited original function in sexual development.

Our results suggest that TALE TF networks represent early-evolving developmental mechanisms. That said, the emergence of complex multicellularity doubtless required more than TF networks. TF-based developmental cues need to be propagated via chromatin-level regulatory mechanisms that establish the cellular memory during embryo development. The extent to which chromatin-level regulatory mechanisms are involved in the development of unicellular organisms is a critical question in elucidating the origins of complex multicellularity.

## Materials and methods

### Strains and culture conditions

Axenic *Micromonas commoda* (RCC299) and *Ostreococcus tauri* (OTH95) were maintained in Keller medium [45] in artificial seawater at room temperature. One hundred mL of a 14- day-old culture was harvested for genomic DNA extraction. *Picocystis salinarum* (CCMP1897) was obtained from the National Center for Marine Algae and Microbiota (NCMA), maintained in L1 medium [46] in artificial sea water, and plated on 1.5% Bactoagar-containing media for single-colony isolation. Genomic DNA of *P. salinarum* was then obtained from a culture derived from one colony.

### Phylogenetic analysis and classification of homeobox genes

Archaeplastida algal TALE homeodomains were collected from the available genomes and transcriptomes listed in S1 Spreadsheet. Details of how TALE sequence was collected is provided in S1A Methods. After excluding nearly identical sequences, a total of 96 sequences together with 18 reference TALE sequences were made into the final homeodomain alignment with 70 unambiguously aligned positions with eight gapped and one constant sites. Details of phylogenetic reconstruction is provided in S1B Methods.

### Bioinformatics analysis

The entire TALE collection was divided into multiple groups representing major clades in the homeodomain tree. Each group was individually analyzed by running MEME4.12 in the motif-discovery mode with default option collecting up to 10 motifs at http://meme-suite.org/ [47]. The search provided multiple non-overlapping motifs, many of which were combined according to previously identified domains such as bipartite KN-A/KN-B, ELK, and HD [14] and independent domain searches against the INTERPRO database (http://www.ebi.ac.uk/interpro/) [48]. All the collected TALE-associated homology domains were aligned to generate HMM motifs, which we used to test if these homology domains are specific to the TALE sequences. All the homology domain information was used to locate any error in gene predictions, and gene models were updated if necessary (Details of the gene model curation is provided in S1C Methods).

### Intron comparison

Introns within the homeodomain were collected and labeled as site numbers of the homeodomain (1-63). If an intron is between two codons it is denoted N/N+1, where N is the last amino acid site number of the preceding exon; introns within a codon are denoted N[n/n+1], where n is one or two for the codon nucleotide position relative to the splice-sites.

### Yeast-two-hybrid analysis

*M. commoda* (affixed with Micco), *O. tauri* (affixed with Ostta), and *P. salinarum* (affixed with Picsa) TALE protein coding sequences were cloned by PCR using primers designed herein (S4 Spreadsheet) from genomic DNAs prepared by the phenol/chloroform extraction and ethanol precipitation method. Micco_62153 and Picsa_04684 contained a single intron, whereas all the other nine genes lacked an intron in the entire open reading frame. For cloning of Micco_62153, we synthesized the middle fragment lacking the intron and ligated them via *XhoI* and *ClaI* sites. For cloning details, see S1D Methods.

## Supporting Information

**S1 Fig. Alignment of TALE homeodomain sequences of the Archae-algal collection**. The 106 sequences were made into an alignment after excluding 20 near identical sequences to reduce redundancy. Animal/amoeba/haptophyte outgroup sequences are included as they share homology with Archaeplastida TALEs outside the homeodomain. The three bars above the sequence numbers show predicted alpha helices. Discarded insertions are noted in red arrowheads.

**S2 Fig. Homology domain alignment of KNOX homogogs**. MEIS class outgroup sequences are included at the bottom. Class label is on the left. KN-A, KN-B, ELK, HOMEOBOX, KN-C1, and KN-C2 domains are labeled on the top. Class groups are labeled by colored bars on the left next to the gene names. Yellow, light green, and green shades in sequences show more than 60%, 80%, or 100% similarity in each column. Gaps between KN-A and KN-B and between KN-B and ELK have been eliminated.

**S3 Fig. GLX class is defined by PBL-Chloro domain**. (A-C) Three alignments are adjusted with inserting gaps for direct comparison among different PBL-Chloro domains. (A) GLX-Chloro class members. (B) GLX-Basal class members. (C) Three Viridiplantae sequences with strong MEINOX homology domain. PBC-homology is shared among the Chlorophyta non-KNOX sequences.

**S4 Fig. Extensive homology of Picsa_04995 and Klefl_00021_0250 to Chocr_41034 indicates their classification as PBX-Red**. MEINOX-homology and PBL-Red domains are indicated by red bars below the alignment.

**S5 Fig. Alignment of Class-B TALE proteins in volvocales**. Short motifs are conserved among all members in this class over the entire length of the sequence.

**S6 Fig. Alignment of Mam-A TALE proteins in mamiellophyceae**. Short motifs (Box1-4) are conserved among all members in this class over the entire length of the sequence. Red reverse triangle at 548-549 shows the truncation position of Micco_Mam-A-tr used in Yeast-two-hybrid analysis.

**S7 Fig. Alignment of Mam-B TALE proteins in mamiellophyceae**. A conserved motif is found between 180-197 amino acids in the alignment. Red reverse triangle at 100-101 shows the truncation position of Ostta_Mam-B-tr used in Yeast-two-hybrid analysis. Homology is restricted to a single homology-A domain ouside the homeodomain.

**S8 Fig. Full-length proteins are necessary for mamiellophyceae non-KNOX TALE proteins to form heterodimers**. Left and Right: Yeast-two-hybrid assays on Ade-/His-/Leu-/Trp-medium. The construct information for the prey conjugated with the GAL4 DNA-binding domain and for the bait conjugated with the GAL4 transcriptional activation domain is given in the table below.

**S9 Fig. TALE interaction network defined by this study using yeast-two-hybrid assays**. (A) Summary diagram for the TALE interaction network. (B) Yeast-two-hybrid assays for the cross-species interaction of TALE proteins. Only one of the possible reciprocal combinations of GAL4 domain conjugations is provided for simplicity. Large X indicates no yeast in the sector. -LTHA: Leu-/Trp-/His-/Ade-medium; -LT: Leu-/Trp-medium.

**S10 Fig. Intron-retention pattern suggests parallel evolution of KNOX and non-KNOX group classes from common duplicated TALE ancestors**. (A) Intron locations collected from 12 TALE classes are shown with arrows. Half arrows indicate cases where not all the class members share the position. White arrows indicate shared positions in at least two different classes, and black arrows indicate class-specific positions. The numbers above the consensus sequence show 60 amino acid positions; the three-amino-acid extension is denoted as ‘abc.’ Row color depicts two alternative domain configurations: purple for MEIS/KNOX types, and navy for PBX/GLX types. Class names are colored according to their phylogenetic groups: green for Viridiplantae, red for Rhodophyta, blue for Glaucophyta, and black for outgroups. The numbers following the class names show how many genes provided the intron information. Of the shared positions, purple triangles on the top mark those shared between MEIS/KNOX and PBX/GLX classes, blue triangles mark those shared between GLX and BELL classes, and red triangles mark those shared between KNOX classes. A notable exception is the KNOX-Red1 class, for which three Rhodophyta clades show different intron locations (44/45, 48[2/3] or 53[2/3]), indicating that the 44/45 intron position can indeed be displaced to 48[2/3] or elsewhere, albeit infrequently. The unique 46/47 intron in the PBX-Glauco (Cyapa_20927) would presumably have resulted from a similar displacement in intron position. (B) Distribution of conserved introns among the TALE homeobox classes. Identified TALE classes are mapped on the Arachaeplastida phylogeny. The 44/45 intron is marked by blue outline and the 48[2/3] intron is marked by red outline. Underlines of the class names indicate the presence of a PBC-homology domain. The Archaeplastida phylogeny is modified from figure 1 of Jackson et al. (2015).

**S11 Fig. ELK-domain alignment.**

**S12 Fig. Identification of MEIS homologs in choanoflagellates.**

**S13 Fig. Identification of PBX homologs in choanoflagellates.**

**S1 Spreadsheet Genomic resources used in this study**. A total of 374 homeobox protein sequences are compiled for this analysis, of which 113 TALE protein sequences are collected. The number of total homeobox proteins and TALE superclass were estimated largely from our homeodomain search described in the materials and methods section. Under Genome annotation, ‘Draft’ indicates a genome without annotation, ‘Trans’ indicates a transcriptome assembly.

**S2 Spreadsheet Archaeplastidal homeobox collection of TALE protein analyzed in this study**. For outgroups, only TALE members that are analyzed in this study are included.

**S3 Spreadsheet KNOX domain homology among KNOX classes**

**S4 Spreadsheet Primers used in this study**

**S5 Spreadsheet Yeast-two-hybrid constructs used in this study**

**S6 Spreadsheet Homeobox profile in Trebouxiophyceae S1 Methods** A. Collecting TALE homeobox protein sequences. B. Phylogenetic reconstruction. C. Homology motif/domain search. D. Intron comparison. E. Cloning of Yeast-two-hybrid constructs.

### S1 Notes

A. Lack of TALE TFs in Trebouxiophyceae. B. Horizontal transfer may explain the presence of Rhodophyta TALE heterodimers in *Picocystis and Klebsormidium* of Viridiplantae.

## Acknowledgements

This work was supported by Discovery Grant 418471-12 from the Natural Sciences and Engineering Research Council (NSERC) (to J.-H.L.), by the Korea CCS R&D Center (KCRC), Korean Ministry of Science, grant no. 2016M1A8A1925345 (to J.-H.L.), NSF-IOS0843119 (to A.Z.W.), GBMF 3788 (to A.Z.W.), and NSF CAREER-1453639 (to E.K.). Postdoctoral support of S.J. was from NSF-IOS0843506 (awarded to Ursula W.Goodenough). We thank Mary Berbee and her laboratory members for valuable comments on the manuscript.

## Author Contributions

### Conceptualization

Sunjoo Joo, Alexandra Z. Worden, Jae-Hyeok Lee

### Data curation

Sunjoo Joo, Ming Hsiu Wang, Gary Lui, Jenny Lee, Sebastian Sudek, Jae-Hyeok Lee

### Formal analysis

Sunjoo Joo, Ming Hsiu Wang, Jae-Hyeok Lee

### Funding acquisition

Sunjoo Joo, Alexandra Z. Worden, Jae-Hyeok Lee

### Investigation

Sunjoo Joo, Ming Hsiu Wang, Jenny Lee, Andrew Barnas, Jae-Hyeok Lee

### Project administration

Alexandra Z. Worden, Jae-Hyeok Lee

### Resources

Sunjoo Joo, Ming Hsiu Wang, Eunsoo Kim, Sebastian Sudek

### Supervision

Sunjoo Joo, Jae-Hyeok Lee

### Writing – original draft

Sunjoo Joo, Ming Hsiu Wang, Jae-Hyeok Lee

### Writing – review & editing

Sunjoo Joo, Eunsoo Kim, Alexandra Z. Worden, Jae-Hyeok Lee

